# Vimentin provides target search efficiency and mechanical resilience for dendritic cell migration

**DOI:** 10.1101/2020.12.18.423401

**Authors:** Luiza Da Cunha Stankevicins, M. Reza Shaebani, Doriane Vesperini, Marta Urbanska, Daniel A. D. Flormann, Emmanuel Terriac, Annica K. B. Gad, Fang Cheng, John E. Eriksson, Franziska Lautenschläger

**Affiliations:** Leibniz-Institute for New Materials, 66123 Saarbrücken, Germany; Department of Theoretical Physics, Saarland University, 66123 Saarbrücken, Germany; Centre for Biophysics, Saarland University, 66123 Saarbrücken, Germany; Department of Experimental Physics, Saarland University, 66123 Saarbrücken, Germany; Biotechnology Centre, Centre for Molecular and Cellular Bioengineering, Technische Universität Dresden, 01307 Dresden, Germany; Weston Park Cancer Centre, Department of Oncology and Metabolism, University of Sheffield, S10 3RX Sheffield, UK; Centro de Química da Madeira, Universidade da Madeira, 9020105 Funchal, Portugal; Cell Biology, Faculty of Science and Engineering, Åbo Akademi University, 20520 Turku, Finland; Turku Bioscience Centre, University of Turku and Åbo Akademi University, 20520 Turku, Finland; School of Pharmaceutical Sciences (Shenzhen), Sun Yat-sen University, 510006 Guangzhou, China

**Keywords:** Vimentin, intermediate filaments, amoeboid cell migration, cell mechanics, cell stiffness, cell viscoelasticity, search efficiency

## Abstract

Dendritic cells use amoeboid migration to pass through confined tissues to reach the lymph nodes, and this homing function is crucial for immune responses. The underlying mechanisms for this type of migration remain unknown. As vimentin intermediate filaments regulate adhesion-dependent migration, we analyzed whether they have a similar effect on amoeboid migration. We show that lack of vimentin impairs amoeboid migration *in vitro* in confined environments, and blocks lymph-node homing in mice *in vivo*. Importantly, we show that vimentin-deficient dendritic cells have a lower coupling factor between cell speed and persistence and reduced target search efficiency (e.g., finding a pathogen, or another cell). These data show that the characteristics of vimentin in its dynamic regulation of cell stiffness and load-bearing, and also elastic capacity, appear to explain the coupling between their migratory ability and search efficiency. Taken together, these data show that vimentin provides the specific mechano-dynamics required for dendritic cell migration and for efficient target searching.

**Summary statement:** Vimentin contributes to the mechanical stiffness of cells required for amoeboid cell migration through confined spaces, and improves cell-search efficiency. Vimentin-deficient cells migrate more slowly and their migration speed is less coupled to persistence compared to control cells.

## Introduction

Together with microfilaments and microtubules, intermediate filaments (IFs) are a major component of the cell cytoskeleton. They share polymer assembly properties and form a net of intracellular filaments that provides the cell with mechanical resistance. They are also involved in the processes of cell signaling, adhesion, and migration. Vimentin is the major IF protein that is constitutively expressed in motile cells of mesenchymal origin. This includes, for example, endothelial cells, leukocytes, connective tissue, and fibroblasts. Vimentin overexpression in cancer cells is often associated with increased cell motility and greater cell malignancy (Ngan et al., 2007; Satelli and Li, 2011).

In a representative experiment by Mendez et al. (Mendez et al., 2010), they injected the vimentin protein into MCF-7 mammary duct carcinoma cells, which resulted in rapid cell polarization and a subsequent increase in cell motility. Thus, vimentin has been implicated in different migratory functions of a number of different adherent cell types, especially in relation to the organization and functionality of actomyosin complexes (Costigliola et al., 2017; Jiu et al., 2015; Jiu et al., 2017).

The role of vimentin in cell migration has been primarily addressed in adherent cells. For example, in fibroblasts, vimentin has been shown to orient traction stress (Costigliola et al., 2017; Guo et al., 2013), which was supported by the demonstration that IFs in general are involved in organization of traction forces (De Pascalis et al., 2018). Furthermore, vimentin is important for leukocyte diapedesis, through its organization of the surface molecules critical for both cell adhesion and the following cell transmigration (Nieminen et al., 2006). However, the role of vimentin in the adhesion-independent migration, also referred to as amoeboid migration, remains poorly understood.

Cells of the immune system migrate in an amoeboid mode of motion, which is fundamentally different to the more commonly studied mesenchymal mode of motion. Mesenchymal migration is characterized by actin stress fibers and focal adhesions that connect the cells to the extracellular matrix (Burridge and Guilluy, 2016). In comparison, amoeboid migration is rapid and is characterized by minimal adhesion, high contractility, and fast shape remodeling (Lämmermann et al., 2008; Reversat et al., 2020). Amoeboid migration depends on friction forces with the environment that can be compared to those used by a person climbing up the inside of a chimney (Hawkins et al., 2009).

The mechanical behavior of migrating immune cells is mainly determined by the cytoskeleton. The cytoskeleton is a complex and dynamic structure that is based on filamentous actin, microtubules, and IFs. This also includes cytoskeletal crosslinkers and molecular motor proteins. In addition, the nucleus provides mechanical support for migrating immune cells (Fletcher and Mullins, 2010). The roles of the microfilaments and microtubules and their control of the mechanical properties of cells have been extensively studied (see (Fletcher and Mullins, 2010) for overview). In contrast, the role of IFs in the control of the mechanical properties of cells is not fully understood yet (Guo et al., 2013; Hu et al., 2019; Janmey et al., 1991; Mendez et al., 2014), and details of the mechanisms behind the involvement of vimentin are still lacking (Danielsson et al., 2018).

Most migration studies have relied on cells that adhere to a two-dimensional (2D) surface (Chernoivanenko et al., 2013; Fletcher and Mullins, 2010; Gan et al., 2016; Lauffenburger and Horwitz, 1996), and only a few studies have addressed the role of vimentin IFs in low-adherence cells. However, studies performed in leukocytes have suggested the importance of vimentin for cytoplasmic rigidity (Brown et al., 2001).

We have previously shown that the mechanical properties between suspended and adherent cells are fundamentally different (Chan et al., 2015). In suspended cells, the cytoskeleton is adapted to the low-adhesive state of the cells, and the cytoskeletal control of cellular mechanics is likely to be different from that of adherent cells. The cytoplasmic localization of vimentin in lymphocytes suggests that vimentin provides mechanical stability to the cell body to enable transmigration through the endothelium (Brown et al., 2001). We hypothesized that the vimentin network governs the dynamical regulation of the stiffness and elasticity of cellular regions, thereby generating the forces that underlie the amoeboid migration of low-adherence immune cells.

Dendritic cells (DCs) have a continuous migratory activity. They are the link between innate and adaptive immune responses, and in their immature stages, they patrol the environment in search of pathogens (Chabaud et al., 2015; Heuzé et al., 2013). DCs are located in peripheral tissues and migrate through highly confined spaces. To do so, DCs have to constantly adapt their shape, which implies a rearrangement of the cytoskeleton. Depending on their function, DCs will adopt diverse searching strategies. On the one hand, immature DCs extensively explore the environment to capture antigens (Moreau et al., 2019). On the other hand, mature DCs are sensitive to hydraulic resistance, so as to find the shortest path toward the lymph nodes where they present the captured antigens to T cells (Chabaud et al., 2015; Renkawitz et al., 2019).

An intermittent random walk (Bénichou et al., 2011) has been described as an alternation between a slow erratic migration and a fast persistent migration. The balance between these two phases allows the exploration of all types of behavior, from a fully diffusive trajectory (i.e., Brownian), to a fully persistent trajectory. We have previously reported that actin waves are involved in immature bone-marrow-derived dendritic cells (BMDCs) switching between Brownian motion and persistent motion (Stankevicins et al., 2020). The role of vimentin in amoeboid cell migration and searching is still poorly understood.

In this study, we aimed to determine whether the lack of vimentin alters immune cell migration through confined spaces. As a model system, we chose primary BMDCs from vimentin wild-type (WT) and vimentin-null (knock-out (KO)) mice, which can migrate through narrow passages, and are required for immune responses. We analyzed immature BMDC migration *in vitro* through measurements of cell speed and persistence in confining one-dimensional (1D) channels and with two-dimensional (2D) plate-plate confiners. By correlating BMDC speed and persistence, we show that these two parameters are less correlated in vimentin-KO BMDCs. We showed previously that such reduced correlation results in less efficient search potential (M. Reza Shaebani, 2020). By quantifying the mechanical properties of BMDCs, we show that vimentin-KO BMDCs are significantly “softer” than WT BMDCs. This suggests that vimentin is indeed required to provide the biomechanical forces that the cell needs for amoeboid migration in confined spaces, and it is therefore also required for efficient search ability. Furthermore, we show that the BMDC *in vivo* homing to lymph nodes is impaired in vimentin-KO BMDCs. These data show that vimentin IFs contribute to the biomechanical properties required for DC migration *in vitro* and *in vivo*, and they suggest a role for vimentin in cell search efficiency of migrating DCs.

## Results

### Vimentin is required for the normal migrating of BMDCs

The confinement of BMDCs has been shown to be crucial for the migration properties of these cells (Heuzé et al., 2013). Therefore, we analyzed their migration *in vitro*, using conditions that mimic the confined physiological environment of BMDCs *in vivo*, as described in previous studies (Le Berre et al., 2014; Liu et al., 2015; Maiuri et al., 2015). To confine the cells in a reproducible manner, and to acquire large datasets of trajectories, we produced 1D confining channels (Fig. 1A, microchannels) and 2D confining plates (Fig. 1B, confining roof), using microfabrication methods (Heuzé et al., 2011; Lautenschläger and Piel, 2013). These devices were then used to analyze the ability of BMDCs to migrate in one dimension and two dimensions.

**Figure 1.**
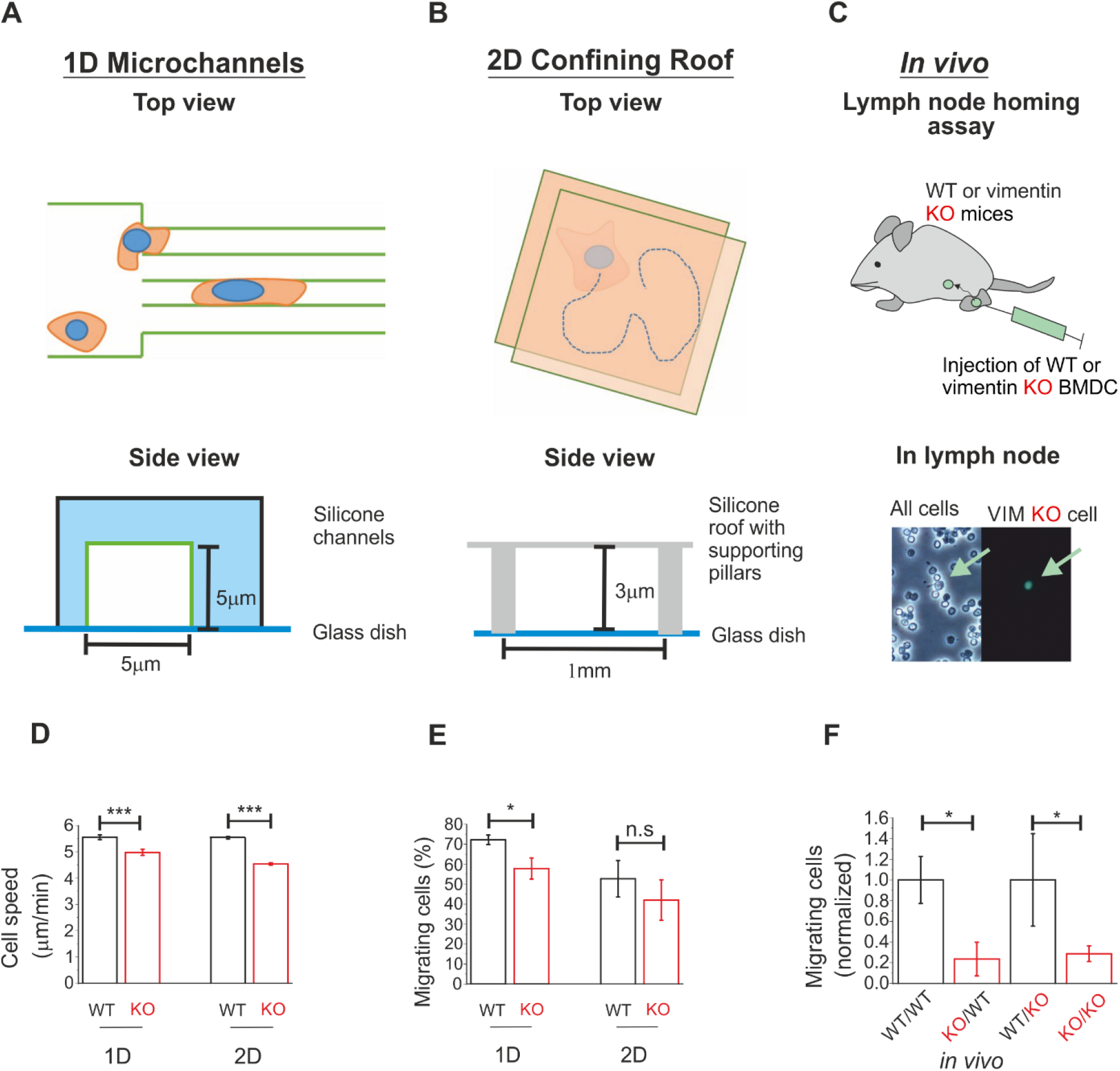
Loss of vimentin results in defective amoeboid migration. (A) Top and side views of one-dimensional (1D) channel migration set-up. (B) Top and side views of two-dimensional (2D) roof set-up. (C) Top: Scheme of lymph node homing assay. Bottom: Representative phase contrast (left) and fluorescence (right) microscopy images used to count the fluorescently labeled vimentin knock-out (KO) cells that arrived at the lymph node. Green arrow, a vimentin-KO cell, from an experiment with vimentin-KO cells injected into the footpad of the mouse. (D) Speed of migrating wild-type (WT) and vimentin-KO BMDCs in one dimension (WT: *n* = 1139 tracks, *v* = 5.55 ±0.09 μm.min^−1^; KO: *n* = 562 tracks, *v* = 4.98 ±0.12 μm.min^−1^; *v*, mean speed ±SE) and two dimensions (WT: *n* = 2428 tracks, *v* = 5.53 ±0.05 μm.min^−1^; KO: *n* = 2409 tracks, *v* = 4.53 ±0.05 μm.min^−1^). (E) Proportion (%) of migrating WT and vimentin-KO BMDCs in one dimension (WT: 72.2% ±2.4%; KO: 57.8% ±5.3 %; mean migrating cells ±SE; 5 independent experiments) and two dimensions (WT: 52.7% ±9.1%; KO: 42.0% ±10.1%; 3 independent experiments). (F) Normalized ratio of injected WT and vimentin-KO BMDCs arriving at a lymph node in WT (WT/WT and KO/WT) or vimentin-KO (WT/KO and KO/KO) mice. \**p* <0.05; \*\*\**p* <0.001; n.s., not significant (linear mixed effects model). Data are means ±SE.

Bone marrow precursors were extracted from wild-type (WT) mice and vimentin knock-out (KO) mice, and following differentiation of the primary DCs, BMDCs were generated (see Methods). Overall, compared to the WT BMDCs, there was significantly lower cell migration with the loss of vimentin in the BMDCs from the vimentin-KO mice (Fig. 1D-F, Table S1). This was seen as lower total numbers of migrating cells (Fig. 1E), as well as lower migration speed (Fig. 1D), and was observed for both the 1D and 2D migration. Taken together, these data indicate that vimentin is essential for these BMDCs to achieve high migration rates and high fractions of migrating cells in 1D or 2D confined environments.

### Vimentin is required for dendritic cell migration to the lymph nodes

We have thus shown here that vimentin is a key component for *in vitro* migration of immature BMDCs. To determine whether vimentin also has a role *in vivo*, we performed lymph node homing assays. This system consists of analyzing whether mature BMDCs lacking vimentin can reach the lymph nodes, as described in previous studies (Faure-Andre et al., 2008). BMDCs become mature when they encounter pathogens. Therefore, we treated these BMDCs with lipopolysaccharide (LPS), which is a membrane component of bacteria. We then injected LPS-treated BMDCs into the footpads of mice, and counted the numbers of BMDCs in the closest lymph node (popliteal lymph node) after 36 h (Fig. 1C). Here, the homing of the vimentin-deficient BMDCs (from the vimentin-KO mice) in the WT mice was compromised when compared to the WT BMDCs (Fig. 1F). Similar data were obtained for lymph node homing in vimentin-KO mice. These data indicate that the observed deficiencies of the vimentin-KO BMDCs are due to the homing of these vimentin-deficient BDMCs, and not to the vimentin levels in the surrounding microenvironment and/or the receiving tissue(s). Taken together, our observations support the hypothesis that vimentin is essential for *in vivo* BMDC homing.

### Vimentin is required for the search efficiency of migrating DCs

There were significantly fewer migrating cells with lower migration speeds *in vitro* comparing the vimentin-KO BMDCs to the WT BMDCs (Fig. 1D, E). This suggests that vimentin-KO BMDCs will explore the space more slowly, which will affect their efficiency in terms of finding randomly distributed targets. As the detection of harmful pathogens is one of the main functions of immature DCs, we investigated the influence of vimentin on the dynamics and search efficiency of these BMDCs. While the speed of these cells will be an influential factor, we also evaluated the other parameters of their motion, as their persistence (Tejedor et al., 2012) and their persistence–speed (*p–v)* coupling (M. Reza Shaebani, 2020), as these might also have crucial roles in cell efficiency to find targets. The instantaneous angular persistence (*p_i_* = cos *θ_i_*) describes the maintenance of the local direction of motion by the cell (Fig. 2A),where *θ_i_* is the turning angle at position *i*. We thus evaluated the instantaneous cell persistence by calculating the directional change at each recorded position *i*.

**Figure 2.**
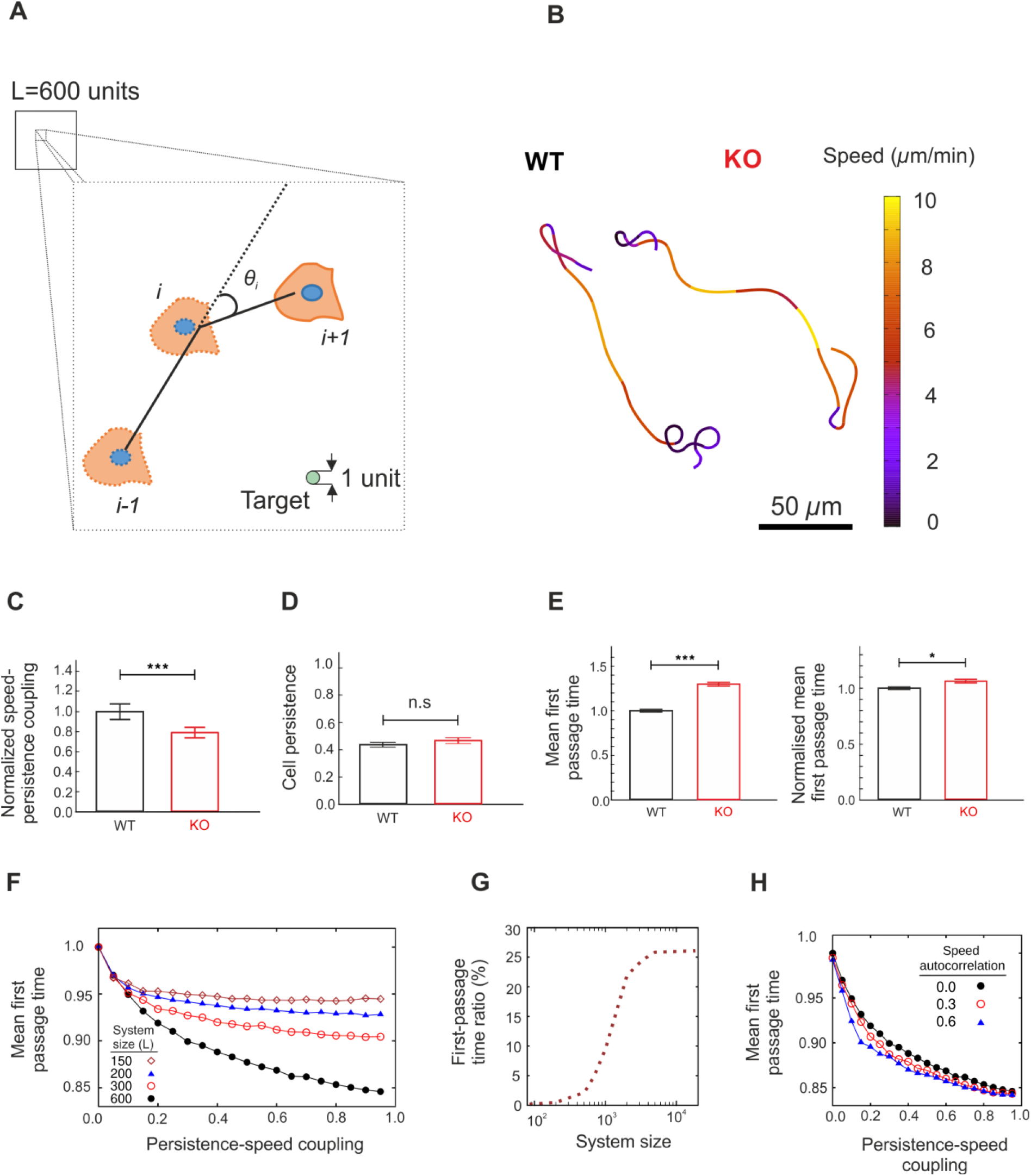
Search efficiency of wild-type and vimentin knock-out BMDCs in two dimensions. (A) Sketch of the migrating area. (B) Sample cell trajectories, color-coded with respect to cell speed. (C) Persistence–speed correlation, *c(p,v)*, scaled by the corresponding value for wildtype (WT) cells. (D) Comparison between mean cell persistence <*p*> of the WT and vimentin knock-out (KO) BMDCs (WT: <*p*> = 0.44 ±0.01, *n* = 3700 tracks; KO: <*p*> = 0.47 ±0.01, *n* = 6500 tracks). (E) Left: Conditional mean first-passage time (MFPT) *τ*, scaled by *τ_WT_*, for a lateral confinement size *L* of 600 (*τ_KO_* = 0.61 *τ_d_*, and *τ_WT_* = 0.47 *τ_d_*, where *τ_d_* is the MFPT for a random Brownian dynamics, with average speed of both cell categories). Right: Similarly, but for MFPT normalized with respect to mean cell speed. (F) Normalized MFPT, scaled by search time for uncorrelated cell persistence and speed 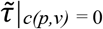, in terms of *c(p,v)* for different system sizes. (G) Relative difference between search time of WT and vimentin-KO BMDC categories 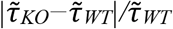 as a function of the system size *L*. (H) MFPT, scaled by search time for uncorrelated motion *c(p,v) = c(v,v)* = 0, in terms of *c(p,v)* for different values of speed autocorrelations. \**p* <0.05; \*\*\**p* <0.001; n.s., not significant (linear mixed effects model).

The turning-angle distributions, P(*θ*), were obtained for both the WT and the vimentin-KO BMDCs (Supplementary Fig. S1A). The resulting probability distributions are qualitatively similar, with mean values around zero (<*θ*> = 0.1° ±1.2°, 0.0° ±0.9°, respectively). When averaged over all of the cells and trajectories (~3700 tracks for WT; ~6500 tracks for vimentin-KO), the mean cell persistence <*p*> were 0.44 ±0.01 and 0.47 ±0.01 for WT and vimentin-KO BMDCs, respectively (Fig. 2D). The persistence of the active random cells is a key factor in the determination of their mean first-passage time (MFPT; average time needed for a cell to reach the position of a target for the first time) (Tejedor et al., 2012). However, we expect that the relatively small difference between the persistence of these WT and vimentin-KO BMDCs (Fig. 2B) will not be sufficient to induce the considerable differences in their MFPTs.

Therefore, we analyzed the coupling between the instantaneous cell speed *v_i_* and the instantaneous angular persistence *p_i_* as an additional influential factor that can influence the search efficiency of these BMDCs. The typical cell trajectories of these WT and vimentin-KO BMDCs showed less curved paths at high speed than at low speed (Fig. 2B). To compare the overall persistence–speed coupling of these WT and vimentin-KO BMDCs, their *p–v* correlation coefficients *c(p,v)* were calculated. The persistence and speed of vimentin-KO BMDCs were significantly less correlated (*c_KO_(p,v)* = 0.69 ±0.04) than for WT BMDCs (*c_WT_(p,v)* = 0.87 ±0.03), as shown in Figure 2C. This difference was 20.7% on average, which indicates that WT and vimentin-KO BMDCs have different migration dynamics. We expect that the combination of these differences between the speed and the *p–v* coupling of these WT and vimentin-KO BMDCs determined their relative search efficiencies.

To better understand the influence of cell migration dynamics on the first-passage properties, we performed Monte Carlo simulations of a correlated active search process by introducing coupling between cell persistence and migration speed (see Methods for details). We chose the target diameter and the searcher diameter as the length unit, as nearly one thirtieth (3.3%) of the mean persistence length of the WT BMDCs. This value was obtained by averaging over the local persistence lengths, *l_p_*, extracted from the instantaneous angular persistence, as *p_i_* = exp(*-d/l_p_*), where *d* is the distance between the recorded positions *i*-1 and *i* (M. Reza Shaebani, 2020). We checked that changing the size of the target or the searcher in our simulations does not influence the reported trends and our conclusions. The speed distributions for WT and vimentin-KO BMDCs were assumed to be uniform in these simulations, for simplicity, with the mean values adjusted to the mean experimental values. The algorithm allowed for the variation of the instantaneous persistence according to the experimental *c(p,v)* value for each category. We defined the conditional MFPT τ as the mean time to reach the hidden target for the first time per unit search area, after excluding the unsuccessful trials (thus, the tail of the first passage time distribution is truncated at the maximum simulation time for each searcher). The conditional MFPT τ was nearly 30% higher for vimentin-KO BMDCs, as shown in Figure 2E. As expected, one reason for the shorter search time of the WT BMDCs is that they migrate faster, where the speed ratio was *v*_WT_/*v*_KO_ ≃ 1.2 (see Fig. 1D). To exclude the contribution of the speed to the MFPT, we introduced a characteristic time *τ\** = 1/*v* to travel a unit of the length with speed *v*. Then, we scale *τ* by *τ*\*, to obtain a normalized MFPT 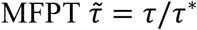 with respect to the mean speed. As shown in Figure 2E, the normalized search time of the WT BMDCs was still significantly lower than the vimentin-KO BMDCs 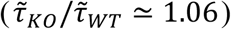. The reason for the remaining difference is most likely the stronger *p–v* coupling for the WT BMDCs. By increasing the coupling strength in simulations, the MFPT monotonically decreased (Fig. 2F). This effect, however, weakens when the confinement size *L* was decreased (i.e., the extent of the environment in which the cell searches for the target). We have previously shown that the *p–v* coupling becomes disadvantageous when the confinement size is smaller than the persistence length of the searcher (M. Reza Shaebani, 2020). The question arises whether in the natural living environments of immature DCs the difference in the *p–v* coupling strength of WT and vimentin-KO DCs is in the range where it significantly affects their mean search time. To answer this, we performed simulations where the persistence *p* and the coupling strength *c(p,v)* of WT and vimentin-KO BMDCs were kept fixed at their experimental values, while the size of the lateral confinement *L* was varied. Figure 2F shows that the relative difference in the normalized search times of WT and vimentin-KO BMDCs, i.e., 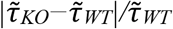 increases with the lateral confinement size *L* and can even exceed 25%. The larger the area a BMDC has to patrol, the stronger the effects of vimentin on cell migration.

In the simulations presented above, we assumed that the successive instantaneous speeds were not correlated. This is the case for a searcher that is free to change its speed after taking each step. The searcher in our simulations “chooses” a new speed at each step from a uniform distribution, for simplicity. However, the analysis of the experimental data showed that there was a weak positive autocorrelation *c(v,v)* between the migration speeds at successive recorded positions (*c(v,v)*, ~0.3 for both WT and vimentin-KO BMDCs, see Supplementary Fig. S1B). This means that the successive speeds at the recorded positions are relatively similar to each other; thus, the cell migration speeds vary slowly with time, and sudden speed changes are rare events.

To see how the speed autocorrelation influenced the MFPT, simulations were performed where the new instantaneous speed *v_i_* remained around the previous speed *v_i-1_* according to the autocorrelation *c(v,v)* values obtained from the experiments. The simulation data presented in Figure 2H confirm that the MFPT was only weakly affected upon variation of the speed autocorrelation, and the trend in terms of the *p–v* coupling strength remained valid, independent of the choice of *c(v,v)*. Additionally, we note that the difference between the speed autocorrelation of the WT and vimentin-KO BMDCs was not significant, as shown in Supplementary Figure S1B; i.e., the loss of vimentin did not result in a significant change of the speed autocorrelation.

Taken together, these data show that while the vimentin-deficient cells retained their relative directional persistence, their persistence–speed coupling was decreased. This lack of correlation between speed and persistence leads to a longer time to find a target (e.g., pathogen), and thus to compromised search efficiency.

### Vimentin is important for the elastic, and not the viscous, properties of BMDCs

We have previously shown that cell mechanics and cell migration are strongly interdependent (Lautenschläger et al., 2009), and vimentin has been reported to control the mechanical properties of cells (Danielsson et al., 2018; Guo et al., 2013). When the subcellular localization of actin and vimentin IFs in immature BMDCs were determined 24 h after their seeding on glass, they showed predominantly cytoplasmic subcellular localization of vimentin filaments, with no detectable vimentin at the cell periphery or in cell protrusions (Supplementary Fig. S2). However, during migration through tissues, the cell body of DCs is confined by other cells and the extracellular matrix (Heuzé et al., 2013; Randolph et al., 2005). Therefore, to determine the subcellular localization of vimentin in a context that better mimics the *in vivo* situation, we imaged cells in custom-engineered 1D-confining channels (Heuzé et al., 2011; Heuzé et al., 2013). Also here, vimentin IFs were predominantly localized close to the nucleus, with no signals at the periphery or in cellular protrusions (Supplementary Fig. S2). These observations are in line with earlier reports that vimentin IFs localize predominantly around the nucleus in cells (see (Danielsson et al., 2018) for overview), and that peripheral cell parts show less prominent IF structures.

As the BMDCs showed predominantly cytoplasmic localization of vimentin IFs (Supplementary Fig. S2), we hypothesized that vimentin mainly controls the mechanical properties of the central, and not the peripheral, cortex of these cells. To test this hypothesis, the mechanical properties of BMDCs were analyzed with and without the expression of vimentin at low and high strain. To measure the cell mechanical properties under low deformation, a high-throughput method was implemented that deformed the cells with a hydrodynamic shear flow on a millisecond timescale, known as real-time deformability cytometry (RT-DC) ((Otto et al., 2015); Fig. 3A). Due to these characteristics of the method, it probes predominantly the outer, cortical region of the cell. The applied shear flow can be adapted to tune the deformation forces imposed on cells. A flow rate of 0.16 μL.s^−1^ (Supplementary Fig. S3, Fr1) was used to cause low deformation of the cells. Only a minor difference was seen for the deformation values between the WT and vimentin-KO BMDCs (Supplementary Fig. S3A-C, Supplementary Table S2).

**Figure 3.**
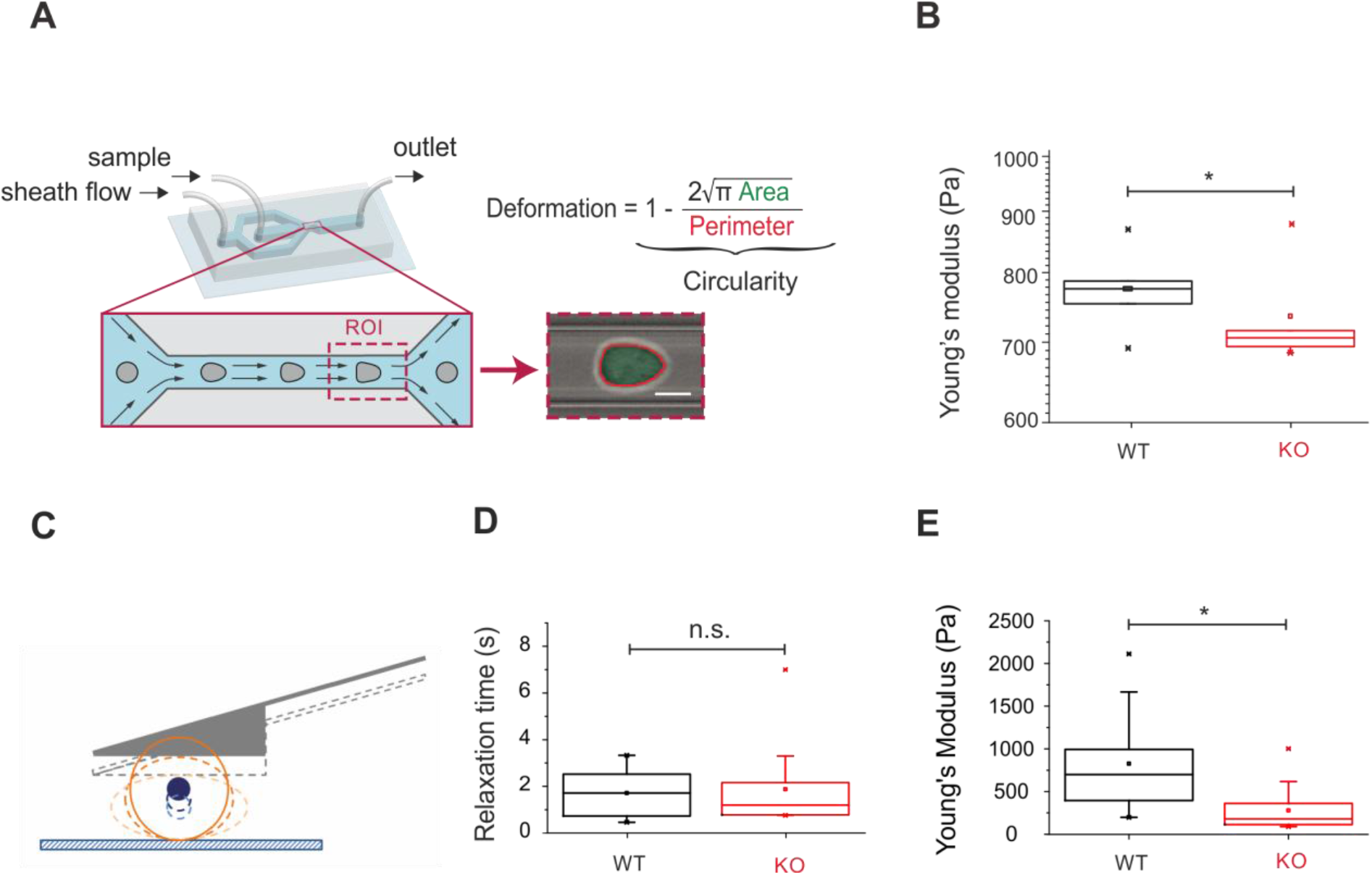
Loss of vimentin in BMDCs decreases cell stiffness. (A) Deformability measurements using real-time deformability cytometry were performed in a microfluidic chip (shown in the background) at the end of a channel constriction (zoom - in). Cell deformation was evaluated using image-derived parameters. (B) Young’s modulus of wild-type (WT; black) and vimentin knock-out (KO; red) BMDCs, for a flow rate 0.16 μL.s^−1^. Box plots of medians from 5 independent experimental replicates; boxes show 25^th^ to 75^th^ percentile range, with a line at the median. Whiskers indicate extreme data points within 1.5× interquartile range (IQR). (C) Scheme of wedged cantilever compressing a cell. (D) Relaxation time of WT and vimentin-KO BMDCs measured at the first extend: results of 3 independent experiments (total 12 WT, 16 vimentin-KO BMDCs). (E) Young’s modulus of WT (black) and vimentin-KO (red) BMDCs measured at the first extend. Whisker middle line represents median values. Means represented by squares. \**p* <0.05; n.s., not significant (linear mixed effects model).

As the force experienced by cells of dissimilar sizes in a microfluidic channel of fixed dimensions is different, it was necessary to extract a material property, such as the Young’s modulus, for any comparative conclusions about the measured mechanical properties of the cells. The Young’s modulus describes the elastic properties of the cell, and expresses how much stress is required to deform a cell by a certain amount. A material with high Young’s modulus is generally described as *stiff*, and one with low Young’s modulus as *soft*. To calculate the Young’s modulus of the deformed BMDCs using RT-DC, we also measured the cell cross-section, where the vimentin-deficient cells were seen to be significantly smaller (Supplementary Fig. S3E). Considering the smaller areas of these cells, the calculated Young’s modulus was significantly lower for the vimentin-KO BMDCs compared to the WT BMDCs (Fig. 3B). This trend was more prominent when the flow rate was doubled (Fr2, 0.32 μL.s^−1^) and the WT and vimentin-KO BMDCs were more deformed (Supplementary Fig. 3C-E).

To clarify the role of vimentin in the determination of the mechanical properties with the higher contribution of the cytoplasmic region of the BMDCs, a low-throughput method was used that offered better control of the speed and depth of cell indentation compared to RT-DC, and that additionally offered viscosity measurements: atomic force microscopy (AFM). For global measurements of their mechanical properties the BMDCs were placed in non-adhesive PEG-coated dishes and compressed using wedged tip-less cantilevers (Fig. 3C; Supplementary Fig. S4A) (Fischer-Friedrich et al., 2016). The cell force relaxations were measured while the cells were compressed. The analysis of the deformation curves, allowed evaluation of the Young’s modulus to describe the cell elasticity, where at an indentation of 4 nN, this was significantly lower in the vimentin-KO BMDCs compared to the WT BMDCs, thus further supporting the data from the RT-DC measurements (Fig. 3E; Supplementary Table S3). This trend persisted also with higher indentation depths, which resulted in stronger deformation (Supplementary Fig. S4C; Supplementary Table S3).

From the analysis of the deformation curves, we also deduced the relaxation time, to determine the cell viscosity, where a higher relaxation time indicates a higher viscosity. AFM is, therefore, ideally suited to describe the viscoelastic properties of cells. The vimentin-KO BMDCs showed no differences in their relaxation time compared to the WT BMDCs (Fig. 3D; Supplementary Fig. S4D; Supplementary Table S3). Hence, as there was a difference in elasticity but not in viscosity, these data suggested that vimentin is important for the elastic properties, and not the viscous properties, of these suspended BMDCs.

## Discussion

Adaptive immune responses rely on rapid, spatiotemporally concerted migration of BMDCs to the lymph nodes or to a site of inflammation (Hampton and Chtanova, 2019). Although many parts of these processes are well understood, little is known about the mechanisms that underlie the actual migration and passage of these cells through complex tissue settings and highly confined spaces. It is clear that the migratory processes involved will require active engagement of the cytoskeleton, as a number of studies have demonstrated specific roles of microtubules and microfilaments (Kopf et al., 2020; Korb et al., 2004). However, less is known about the functions of IFs, although vimentin has been demonstrated to have an essential role in leukocyte homing (Nieminen et al., 2006). Therefore, we wished to specifically determine the contribution of vimentin to the biomechanics required for BMDC migration.

To investigate the role of vimentin in the migration of immune cells, we studied confined, amoeboid migration using 1D and 2D techniques that allowed for long (15 h) periods of observation. While migration in two dimensions provides additional parameters, such as the migration path length and the angular persistence, the 1D experiments enforced confinement all around the sides of the cell, which might be more reflective of the local geometries that they encounter during migration through very tight spaces. We observed that the migration speeds of the vimentin-KO BMDCs were lower than the WT BMDCs for both one dimension and two dimensions. This finding suggested that vimentin is involved in the motility of these cells, as has been shown for mesenchymal cell migration (Gan et al., 2016; Patteson et al., 2019a).

In our previous studies, we showed that there is a universal coupling between cell speed and cell persistence (Maiuri et al., 2015; Wu et al., 2014); i.e., these two quantities of migration are not completely independent variables. In particular, following our analysis of a range of different cell types, we revealed a relatively strong general trend that faster cells migrate straighter compared to slower cells (Maiuri et al., 2015). This correlation remained true within a cell type population where faster elements migrate in a more persistent manner. We have also previously shown that high *p–v* coupling reduces the search time of migrating cells (M. Reza Shaebani, 2020). In our 2D experiments, vimentin-KO BMDCs showed a lower coupling coefficient between speed and persistence, *c(p,v)*, compared to WT BMDCs. This suggested that they will take more time to find randomly distributed targets (e.g., pathogens). The search efficiency of these WT BMDCs should be visibly greater than vimentin-KO BMDCs, not only because they are faster, but also because their speed and persistence are more strongly coupled.

Let us consider dermal DCs in the skin or DCs in small intrapulmonary airways for comparison: the cell density ranges from <100 to a few hundred cells per mm^2^ in these environments (Ng et al., 2008; Schon-Hegrad et al., 1991). Assuming a density of 50 cells.mm^−2^, the natural searching environment per cell is 0.02 mm^2^. In such a case, each cell patrols an area corresponding, for example, to a square with 140 μm sides, which is roughly 14-fold the typical cell persistent length, which is in the range of 10 μm. This corresponds to a system size of greater than 600 in our simulations. At such system sizes, the lower *p–v* coupling strength of the vimentin-KO BMDCs will trigger nearly 10% longer search times (according to Fig. 2F). Additionally, the lower speed of the vimentin-KO BMDCs will further lower their search efficiency by ~20 %.

In a previous study, we showed that cell mechanics and immune cell migration are coupled (Lautenschläger et al., 2009). Therefore, we tested here whether the decreased migratory abilities of these vimentin-KO BMDCs are due to a change in their mechanical properties. By comparing two kinds of mechanical measurements, we showed that the lack of vimentin affects the elasticity of these cells at both low (i.e., using RT-DC) and high (i.e., using plate-plate AFM) strain without affecting their viscosity. The contribution of polymers with different mechanical properties can result in strong networks (Janmey et al., 1991). *In vitro* rheology analysis of the cytoskeleton filaments have revealed that although vimentin is more deformable compared to actin or microtubules, it is more resistant to breakage at high strain and provides greater toughness to living cells (Hu et al., 2019; Janmey et al., 1991). Our observation here that this loss of vimentin in suspended BMDCs results in softer cells is consistent with previous observations in adherent cells (as we reviewed in (Danielsson et al., 2018), and was recently shown in (Patteson et al., 2019b)). Furthermore, our findings show that vimentin provides elasticity to these suspended BMDCs, which has a role when cells need to deform or squeeze through confined environments.

This question was also investigated in a recent study on mesenchymal fibroblasts, where the hyperelastic properties of vimentin were measured (Hu et al., 2019). This study showed that vimentin-deficient cells have a lower force relaxation than WT cells, and vimentin-enriched cells have higher force relaxation than WT cells. This suggests that vimentin is a hyperelastic fiber; i.e., a fiber that responds elastically even under large deformation. This indirectly contributes to cell viscoelasticity and mesenchymal cell migration (Guo et al., 2013; Hu et al., 2019; Smoler et al., 2020). In the light of previous knowledge, our data indicate that when cells migrate in an amoeboid fashion, they can use both the load-bearing and the elastic properties of vimentin to generate the required dynamic forces for this type of motility. As vimentin organization and polymerization is dynamically regulated by constitutive phosphorylation and dephosphorylation cycles, this will provide the dynamic properties needed for meaningful contributions to the required forces (Eriksson et al., 2004; Helfand et al., 2011; Ivaska et al., 2007). The differences between migration behavior of mesenchymal and amoeboid migrating cells under the loss of vimentin can be seen by comparison of our data to a recent study that investigated the migration speeds in one dimension and two dimensions of fibroblasts that lacked vimentin (Patteson et al., 2019a). Interestingly, these fibroblasts showed similar migration behaviors to the BMDCs for the 2D surfaces (i.e., slightly lower migration speed), but an opposite effects in the 1D microchannels (i.e., increased migration speed of fibroblasts) (Patteson et al., 2019a). This emphasizes the differences between these two migration types, which become particularly evident in one dimension. Moreover, it has been shown in adherent cells, that vimentin provides localized increases in cell stiffness and protects the nucleus from rupture (Patteson et al., 2019b). For this reason, vimentin-deficient cells can show stronger deformation of the nucleus itself in confined spaces, which makes the mesenchymal cells accelerate in the 1D microchannels (Patteson et al., 2019a). Considering that vimentin filaments in migrating DCs are localized mainly in the cytoplasm, we suggest that vimentin reinforces this subcellular regions against strong compressive stress. This idea was supported by Patteson et al. (Patteson et al., 2019b), who showed that the absence of vimentin results in an unstable nuclear shape and increased DNA damage during migration through a 3D collagen mesh.

If vimentin protects against DNA damage, and nuclear deformation and instability (Patteson et al., 2019b; Sarria et al., 1994), we can assume a role for vimentin in the modulation of gene expression. In this framework, the lack of vimentin can have broader consequences, as DC differentiation, maturation, and migration rely on changes in gene expression and differential expression of surface receptors. In the present study we explored the mechanical properties of the cytoskeleton. We showed that the interplay between vimentin and actin has a strong effect on the mechanical properties of the cells, while the absence of these interactions might have other consequences.

Taken altogether, our data demonstrate that vimentin contributes to the build-up of the mechanical properties that cells need for amoeboid migration. All of these effects facilitate the migratory abilities of DCs and reduce their MFPT during their search for pathogens.

## Materials and Methods

### Cells

Primary DCs were differentiated from bone-marrow precursors extracted from vimentin WT and vimentin-KO mice. BMDCs were generated as previously described (Vargas et al., 2016), using Iscove’s modified Dulbecco’s medium (IMDM) with 10% fetal calf serum (FCS), 2 mM glutamine, 100 U.mL^−1^ penicillin, 100 μg.mL^−1^ streptomycin, 50 μM 2-mercaptoethanol, and 50 ng.mL^−1^ granulocyte macrophage colony-stimulating factor, and containing supernatant obtained from transfected J558 cells. The semi-adherent cell fraction, which corresponds to the CD86^+^ cells, was gently flushed from the cell culture dishes. All of the experiments were performed during cell differentiation days 10-12. All of the reagents used for cell culture were from Thermo Fisher (Waltham, MA, USA).

### Mice

Vimentin heterozygous mice (129/Sv × C57BL/6) were used to generate vimentin-deficient (KO) homozygotes (VIM^−/-^) and WT offspring. Both vimentin WT and vimentin-KO mice used for the bone marrow extraction were maintained in the Animal Facility of Biocity (Turku, Finland) under permit *7284/04.10.03/2012* and protocol number *197/04.10.07/2013* of the Ethical Committee for Animal Experiments of the University of Turku. The regular molecular cloning and other GMO1-related research work was performed under the permit of the Board of Gene Technology, Finland (GTLK), with notification number *038/M/2007*.

### Antibodies and chemicals

The antibodies used for immunostaining were an anti-vimentin monoclonal antibody (D21H3; Cell Signaling Technology, Danvers, MA, USA) combined with Alexa 647 conjugated secondary antibodies (Abcam, Cambridge, UK). For actin staining, we used phalloidin conjugated with tetramethyl rhodamine B isothiocyanate (Sigma Aldrich, St Louis, MO, USA), and for nuclei staining, Hoechst 34580 (Sigma Aldrich, St Louis, MO, USA). The antibodies used for Western blotting, were the anti-vimentin monoclonal antibody (D21H3; Cell Signaling Technology, Danvers, MA, USA), combined with the secondary anti-horse radish peroxidase (HRP) anti-rabbit antibody (Bio-Rad, Hercules, CA, USA), and an anti-biotin monoclonal antibody (Hsc70; 1B5; Enzo Life Sciences, Plymouth, PA, USA).

### Channels

The microchannels used in the 1D migration experiments were manufactured according to previously published procedures (Heuzé et al., 2011; Vargas et al., 2014). Briefly, silicone rubber kits (RTV-615; Momentive Performance Materials, USA) were used to mix the silicon and the curing agent (10:1), which was then degassed and polymerized at 75 °C in the specific microfabricated molds of the positive channel imprint. The resulting 1D channels of 5 μm height and 5 μm width were sealed in 35-mm glass bottomed cell culture dishes (World Precision Instruments, Sarasota, FL, USA) using plasma surface activation. The assembled structure was coated with 20 μg.mL^−1^ fibronectin (Sigma-Aldrich, St Louis, MO, USA) and incubated with cell culture medium for 60 min. The cells were platted at the channel entry at a concentration of 2 ×10^7^ cells.mL^−1^.

### Plate-plate confiners

The cell-confining plate “roofs” were prepared as previously described (Le Berre et al., 2014; Liu et al., 2015). The mold for the coverslip poly(dimethylsiloxane) (PDMS) coating was produced by photolithography, and consists of pillars of 3-μm height and 440-μm diameter, spaced at 1000 μm from each other. The confiner was assembled in glass-bottomed six-well plates (Mattek, Ashland, MA, USA). The roof of the 3-μm height spacers was covered by a 10-mm coverslip. The roof was adhered to the culture dish lid using a softer, deformable PDMS pillar that was later closed and fixed with adhesive tape. Before the experiments, both the cell culture dishes and the confiner roofs were coated with 0.5 mg.mL^−1^ poly(L-lysine)-graft-poly(ethylene glycol) (PLL-PEG) (SuSoS, Dübendorf, Switzerland) to avoid cell adhesion. The mounted device was incubated in culture medium prior to the experiment, for at least 4 h, to equilibrate the PDMS. Cell recording started 2 h after the cell confinement.

### Migration assays cell trajectories

Cell nuclei were stained with 200 ng.mL^−1^ Hoechst 34580 (Sigma Aldrich, St Louis, USA), for 30 min, and cell migration was recorded using epifluorescence microscopy, over at least 6 h. The cells were kept at constant atmosphere of 37 °C and 5% CO2 (Okolab, Italy) during the entire experiment. Images were obtained using the inverted microscope system (eclipse Ti-E; Nikon, Tokyo, Japan) equipped with a fluorescent illumination. Cell trajectories were tracked using the custom-made software described by (Maiuri et al., 2012). For the analysis, all of the trajectories were filtered to exclude those <25 μm and <30 min. The number of migrating cells was then measured as the number of remaining tracks, according to these thresholds. Migrating cells that passed through the acquisition field for <30 min, were then not included in the analysis.

### In vivo *lymph node homing assay*

The *in vivo* migration assay was performed according to (Lämmermann et al., 2009), with some modifications. Prior to injection of the mice, BMDCs from WT and vimentin-KO mice were treated with 100 ng.mL^−1^ LPS for 30 min, and fluorescently labeled with a cell tracer (Oregon 488; Thermo Fisher Scientific, Waltham, MA, USA), according to the manufacturer protocol. A total of 1 ×10^6^ fluorescently tagged cells diluted in phosphate-buffered saline (PBS) to a final volume of 40 μL were injected subcutaneously into the hind footpad of mice aged between 6 and 10 weeks old. The mice were killed 36 h after BMDC injection, and the popliteal lymph node was extracted. The lymph nodes were then mechanically disrupted and digested using collagenase D (Roche, Basel, Switzerland) and DNAse1 (Roche, Basel, Switzerland) for 30 min at 37 °C. The homogenate was filtered through a 77-μm silk filter. The purified cells were mounted on a microscope slide, and the ratio between the fluorescent and non-tagged cells were counted. This ratio was normalized to the ratio of WT cells that had arrived in the lymph nodes.

### Immunofluorescence

The cells were fixed for 10 min with 4% paraformaldehyde (Sigma Aldrich, St Louis, MO, USA), and permeabilized with 0.5% Triton X-100 (Sigma Aldrich, St Louis, MO, USA) for 10 min. Protein blocking was achieved by incubating the cells with 3% bovine serum albumin (BSA) for 1 h. Primary antibodies were diluted 1:200 in BSA 3 % and incubated overnight at 4°C. Secondary antibodies were diluted at 1:1000 ratio in 3% BSA and incubated for 1 h at room temperature. The actin staining used phalloidin conjugated with rhodamine (1 mM final).

### SDS-PAGE and Western blotting

Whole cell lysates were extracted using Laemmli sample buffer. Protein separation was carried out by SDS-PAGE in 10% bis-acrylamide gels and transferred to nitrocellulose membranes (Amersham Protran; pore size, 0.45 μm; GE Healthcare, Chicago, IL, USA). The membranes were blocked with 5% fat free milk in Tris-buffered saline (TBS) and incubated overnight at 4 °C with the anti-vimentin antibody (dilution 1:1000 in 5% BSA) or with the loading control HSC70 (dilution 1:1000 in 5% BSA). Proteins were detected using enhanced chemiluminescence (Western Lightning Plus-ECL; Perkin Elmer, Waltham, MA). The antibodies used in the assay are listed in the antibody section.

### Confocal microscopy

Confocal images were obtained using a spinning disk unit (CSU W1; Yokogawa; Andor Technology, Belfast, UK) with a pinhole size of 50 μm coupled to the inverted microscope system (Eclipse Ti-E; Nikon, Tokyo, Japan). Images were recorded using a flash 4.0 camera with a 6.5-μm pixel size (Hamamatsu, Hamamatsu City, Japan). Image treatment and Z maximum projection were carried out using the ImageJ software (FIJI) (Schindelin et al., 2012).

### Atomic force microscopy

To measure the global stiffness of the cells in suspension, tipless cantilevers (MLCT-010, cantilever F) with a spring constant of 0.3-1.2 (nominal: 0.6) N m^−1^. (Bruker, Billerica, MA, US) were wedged to correct the 10° cantilever tilt, which provided a flat surface to probe the cells. The cantilevers were wedged according to (Fischer-Friedrich et al., 2016). Tipless cantilevers were pressed against drops of adhesive (Norland Optical Adhesive 63; NOA63; Thorlabs, Newton, NJ, US) placed on a silicon-coated coverslip. The NOA63 was cured with UV light for 60 s, gently detached from the silicon coverslip, and cured for an additional 300 s. The flatness and integrity of the wedged cantilevers was assessed using electron microscopy.

Atomic force microscopy measurements were carried out using JPK Cell Hesion 200 (JPK Instruments, Berlin, Germany) mounted on the inverted microscope system (Eclipse Ti-E; Nikon, Tokyo, Japan). The sensitivity was calibrated by acquiring a force curve on the dish surface, and the spring constant was calibrated by the thermal noise fluctuation method, using the build-in JPK software. For measurements, the cantilever was lowered at a speed of 0.5 μm.s^−1^ with the point force set at 4 nN. Measurements consisted of four subsequent compressions of 1 μm for 60 s each. During compressions, the height was kept constant and the cell force was recorded for all extends (total time, 300 s). Bright-field images were taken for all of the steps of the measurements (magnify 40× plus 1.5× zoom, to calculate the size and contact area of the cells without cellular protrusions). The cells were kept in at 37 °C during the measurements, and the culture media was supplemented with 200 mg.mL^−1^ Hepes buffer. The Young’s modulus was calculated using the Hertz-approximation on each extend (Chugh et al., 2017). The Hertz-model is a purely elastic model, which means that the “Young’s modulus” here was basically an apparent elastic modulus, because cells are viscoelastic objects.

### Real-time deformability cytometry

High-throughput measurements of cell mechanics were performed using RT-DC, according to previously described procedures (Urbanska et al., 2018). In brief, the cells were suspended in 100 μL viscosity-adjusted measurement buffer (0.5% [w/v] methylcellulose [4000 cPs, Alfa Aesar, Germany] in phosphate-buffered saline without Mg^2+^ and Ca^2+^; final viscosity, 15 mPa.s^−1^) and flushed through a 300-μm long microfluidic channel with a 30 × 30 μm square cross-section, at flow rates of 0.16 μL.s^−1^ (Fr1) and 0.32 μL.s^−1^ (Fr2). Images of the deformed cells were acquired at 2000 frames.s^−1^ within a region of interest close to the channel end. Cell deformation and cross-sectional area were evaluated in real-time based on contours fitted to the cells by an in-house developed image processing algorithm (Otto et al., 2015). The data obtained were filtered for cell areas of 50 μm2 to 500 μm^2^, and an area ratio of 1.00-1.05. The area ratio is defined as the ratio between the area enclosed by the convex hull of the contour, and the area enclosed by the raw contour, with the cells with rough or incompletely fitted contours discarded. Young’s modulus values were assigned to each cell using a data grid that was obtained from numerical simulations for an elastic solid (Mokbel et al., 2017), with the aid of the ShapeOut analysis software, version 0.8.7 (available at https://github.com/ZELLMECHANIK-DRESDEN/ShapeOut). A minimum of 500 cells were analyzed per condition in each experiment, with five experiments carried out. Statistical analysis was performed using linear mixed effect models, as described in detail elsewhere (Herbig et al., 2018).

### Simulation method

We performed Monte Carlo simulations of a discrete time persistent random walk process in a square box of size *L* with periodic boundary conditions. We used the experimental tracks as input for simulations. Their initial starting point and orientation was however randomized to use the same trajectory when restarting the search process, with a new imaginary target position. We report the results for the system size *L* = 600 (unless mentioned otherwise). A single random target was considered in each simulation, which is equivalent to regularly spaced targets on an infinite plane with 1/*L*^2^ density. The searcher moves *v* steps at each time step drawn from a speed distribution *P(v)*, in case of uncorrelated successive migration speeds. To incorporate the speed autocorrelation, the sum-of-uniforms algorithm was implemented (Chen, 2005; Lakhan, 1981; Willemain and Desautels, 1993). This algorithm was also used to correlate the cell speed and persistence. The angular persistence *p* varied from 1, for continuing along the previous direction of motion (i.e., *θ* = 0), to −1 for reversing the direction of motion (*θ* = ±π). For a completely random turning (i.e., *θ ε* [-π, π])*p ε* [-π, π] with a minimum absolute value of 0 (i.e., 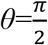 or 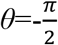). *θ_i_* is the turning angle at position *i*; i.e., the angle between the arrival directions at successive positions *i* and *i+1* (Fig. 2A). The method allows for tuning the desired degree of stochasticity for the resulting speed and persistence. This is achieved by means of an additional parameter Δ. A new speed is generated at each time step from the speed distribution centered around the current speed. The random speed generation via the sum-of-uniforms procedure produces the demanded speed autocorrelation. Next, a new cell persistence was taken from a uniform distribution around the value determined by the *p–v* coupling strength and the instantaneous speed *v*. The algorithm allows for variation of instantaneous persistence in such a way that the overall *p–v* coupling was the experimental *c(p,v)* value for each category. Here the parameter Δ tunes the extent of stochasticity; i.e., the variation range of the cell persistence for the given cell speed. After determining the new *v* and *p*, the searcher moves *v* sub-steps, while it is allowed to vary its orientation after each sub-step according to the current cell persistence.

## Author contributions

L.S., E.T., J.E., and F.L. conceived the experiments. L.S., M.U., D.F., E.T. and F.C. performed experiments. L.S., M.U. and M.R.S. analyzed data. M.R.S. performed numerical simulations. L.S., D.V., M.R.S., A.K.B.G and E.T. wrote the manuscript. L.S., D.V. and M.R.S. drafted the figures. J.E. and F.L. revised the manuscript.

## Acknowledgements

The authors would like to thank K. Kaub, Z. Mostajeran, M. Piel and P. Vargas for helpful discussions concerning BMDC migration and mechanics; and Elisabeth Fischer-Friedrich for advice and discussions on using a wedged cantilever for AFM measurements.

## Abbreviations

AFM: Atomic force microscopy
BMDC: Bone marrow derived dendritic cell
DC: Dendritic cell
IF: Intermediate filament
IMDM: Iscove’s modified Dulbecco’s medium
KO: Knock-out
LPS: Lipopolysaccharide
MFPT: Mean first-passage time
PDMS: Poly(dimethylsiloxane)
PLL-PEG: Poly(L-lysine)-graft-poly(ethylene glycol)
RT-DC: Real time deformability cytometry

## Supplementary Material

**Figure S1.**
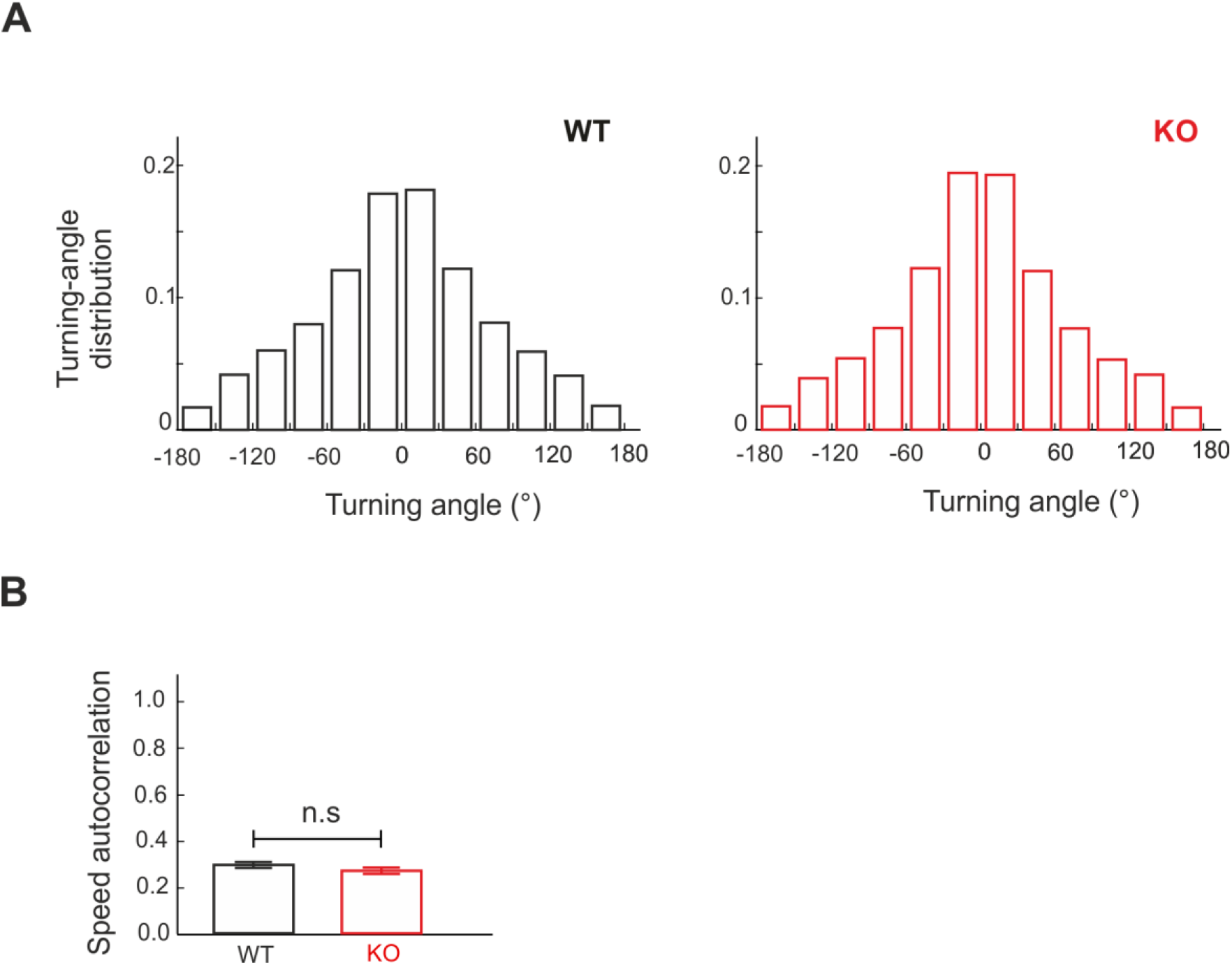
Comparison between directional and speed correlations of wild-type and vimentin knock-out BMDCs in two dimensions. (A) Probability distribution of the turning angle *θ* of wild-type (WT, left) and vimentin knock-out (KO, right) BMDCs at recorded positions. (B) Comparison between the mean speed autocorrelations *c(v,v)* of WT and vimentin-KO BMDCs. n.s., not significant (linear mixed effects model).

**Figure S2.**
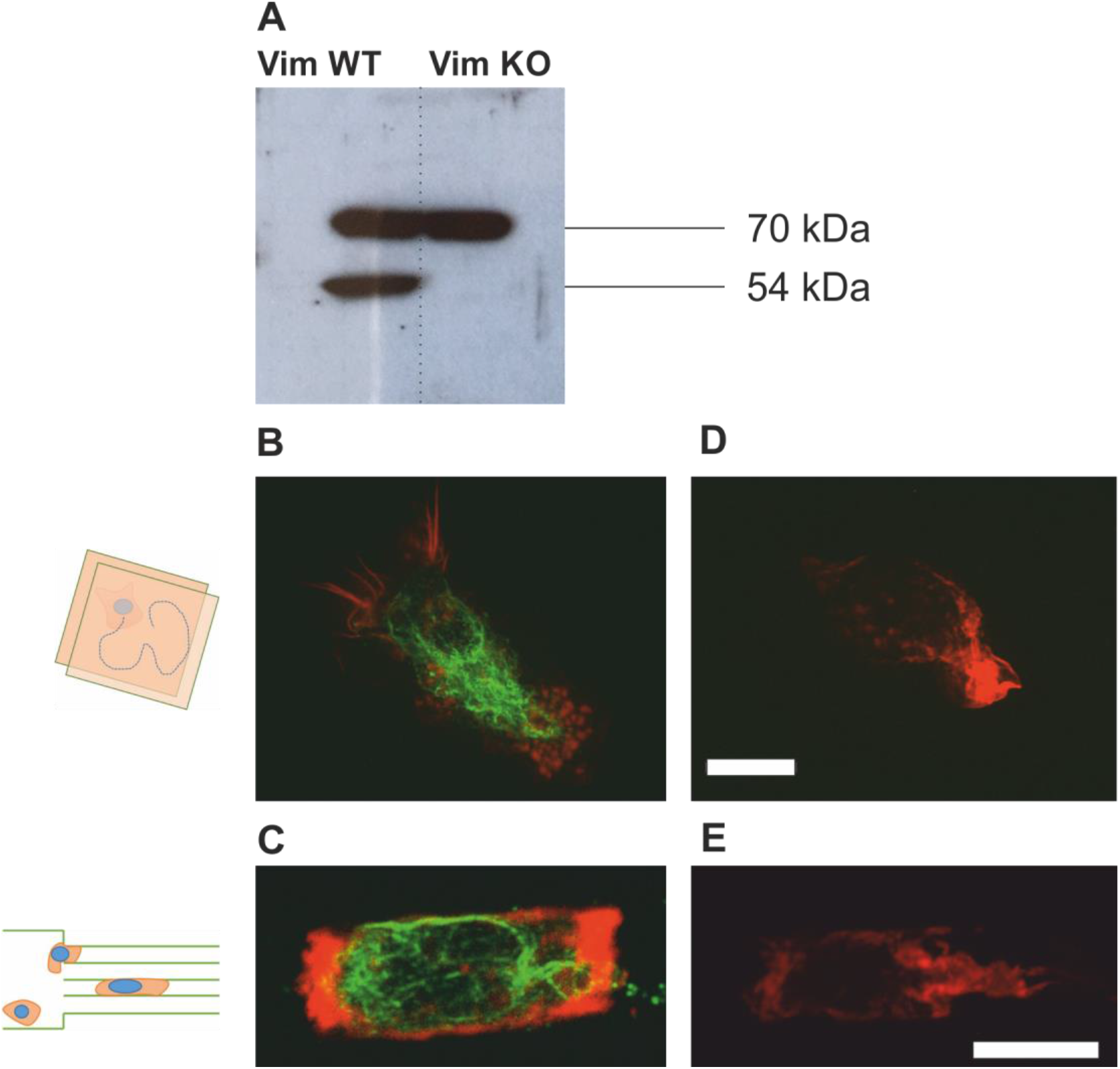
Cytoskeleton observations of WT and vimentin-KO BMDCs in one dimension and two dimensions. (A) Representative Western blot for protein quantification of vimentin in wild-type (WT) and vimentin knock-out (KO) primary BMDCs (54 kDa, vimentin). Loading control Hsc70 (70 kDa). (B-E) Representative images of WT (B-C) vimentin-KO (D-E) BMDCs for actin (red) and vimentin (green). Cells were on a glass coverslip and in 5 μm high and 5 μm wide micro-channels, with all surfaces coated with fibronectin. Scale bars, 10 μm.

**Figure S3.**
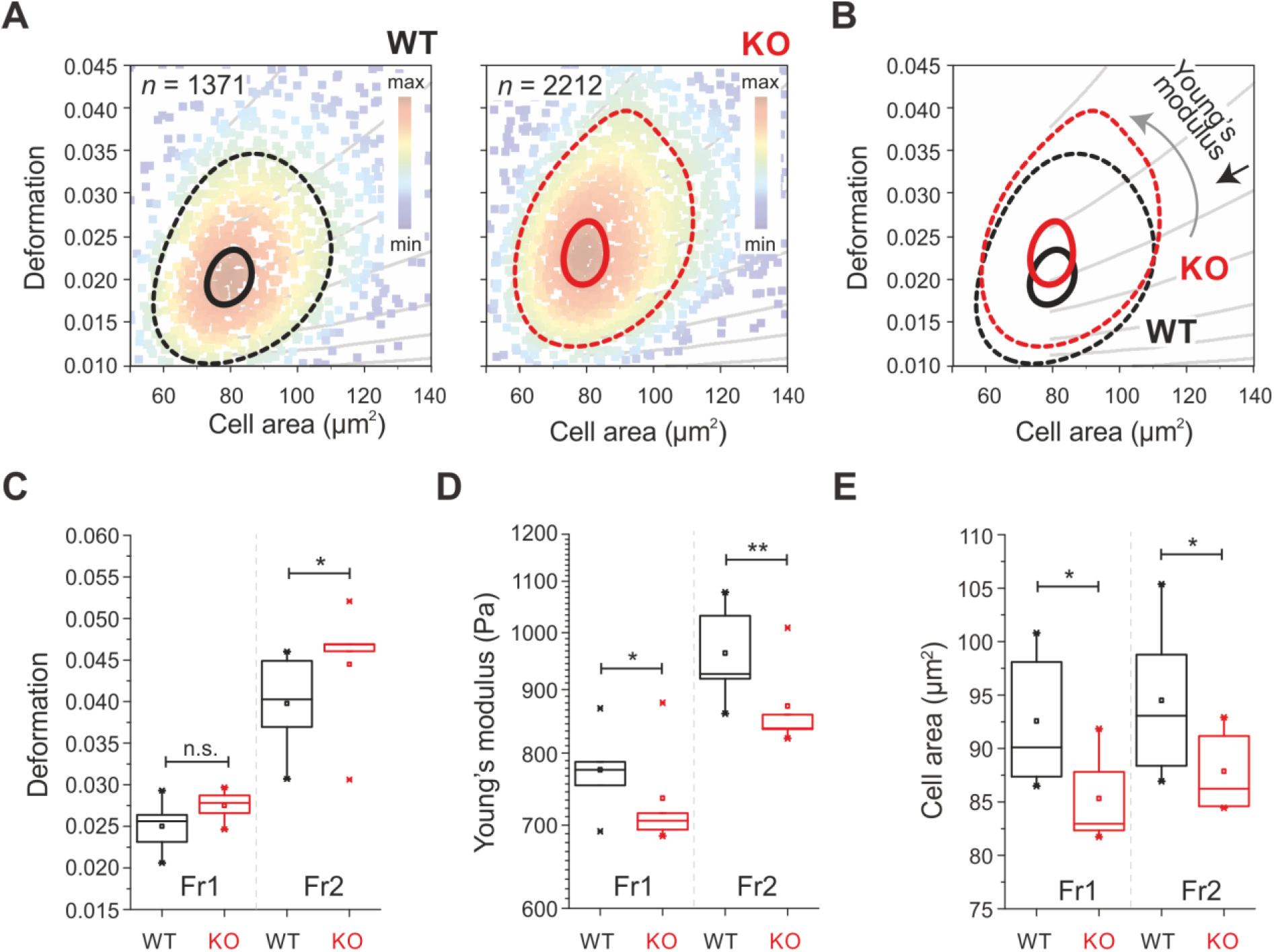
Real-time deformability cytometry analysis of global mechanical properties of wild-type and vimentin-KO BMDCs. (A) Deformation–cell area scatter plots, showing a representative measurements of wild type (WT) and vimentin knock-out (KO) BMDCs. Cell numbers, *n*, as indicated. Color map: event density. Contour plots delineate 50% density (dashed lines) and 95% density (solid lines). (B) Overlay of contours from (A). Grey lines, isoelastic regions from numerical simulations, grouping cells of same mechanical properties. (C-E) Comparisons of deformation (C), Young’s Modulus (D), and cell area (E) of WT and vimentin-KO BMDCs, measured at two different flow rates: Fr1, 0.16 μL.s^−1^; Fr2, 0.32 μL.s^−1^. Box plots of medians from 5 independent experimental replicates shown; boxes show 25^th^ to 75^th^ percentile range, with a line at the median. Whiskers indicate extreme data points within 1.5× interquartile range (IQR). \**p* <0.05; \*\**p* <0.01; n.s., not significant (linear mixed effects model).

**Figure S4.**
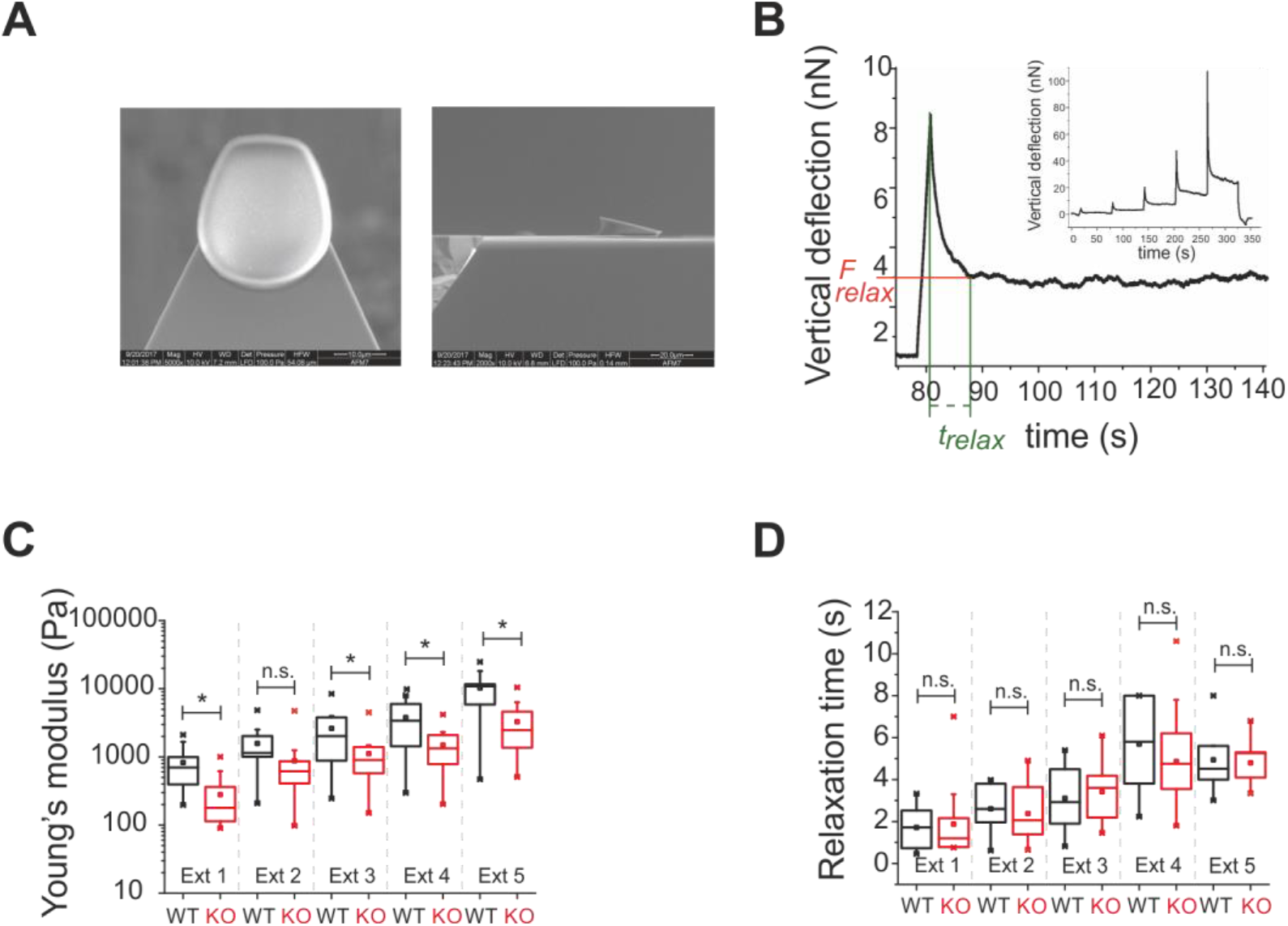
Force-mode atomic force microscopy analysis of dendritic cell mechanics. (A) Representative ventral and lateral electron microscopy images of wedged cantilevers used to evaluate the cell global mechanical response by atomic force microscopy. (B) Scheme of deformation graph used for analysis, showing measurement of relaxation time with atomic force microscopy. (C, D) Young’s modulus (C) and relaxation time (D) of wild-type (WT; black) and vimentin knock-out (KO; red) BMDCs measured at different extends. Data from 3 independent experiments (total: 12 WT, 16 vimentin-KO BMDCs). \**p* <0.05; n.s., not significant (linear mixed effects model). Whisker middle lines represent median values. Boxplot: mean values shown by squares.

**Table S1.** Loss of vimentin results in defective amoeboid migration.

| System | Observable | Cell type | Datum (mean ± SE) |
| --- | --- | --- | --- |
| 1D | Migrating cells (%) | WT | 72.2 ±2.4 |
| | | KO | 57.8 ±5.3 |
| | Migration velocity ($\mu\text{m}.\text{min}^{-1}$) | WT | 5.55 ±0.09 |
| | | KO | 4.98 ±0.11 |
| | Apparent persistence | WT | 0.68 ±0.007 |
| | | KO | 0.66 ±0.01 |
| 2D | Migrating cells (%) | WT | 52.7 ±9.14 |
| | | KO | 42.0 ±10.15 |
| | Migration velocity ($\mu\text{m}.\text{min}^{-1}$) | WT | 5.5 ±0.05 |
| | | KO | 4.5 ±0.05 |
| | Path length ($\mu\text{m}$) | WT | 383.0 ±23.6 |
| | | KO | 240 ±19 |
| | Apparent persistence | WT | 0.51 ±0.004 |
| | | KO | 0.51 ±0.004 |
| *In vivo* | Cells arriving at the lymph node<br>(normalized) | WT/WT | 0.54 ±0.23 |
| | | KO/WT | 0.55 ±0.16 |
| | | WT/KO | 1.00 ±0.50 |
| | | KO/KO | 0.24 ±0.07 |

WT, wild-type; KO, vimentin knock-out

**Table S2.** Loss of vimentin in dendritic cells decreases cell stiffness, as shown by real-time deformability cytometry.

| Flow rate | Cell type | Cells per experiment (n) | Total cells (n) | Deformation | Young's modulus (Pa) | Cell area ($\mu\text{m}^2$) |
| --- | --- | --- | --- | --- | --- | --- |
| 1 | WT | 3,069 | 10,238 | $0.0256 \pm 0.0025$ | $776 \pm 22$ | $90.1 \pm 3.6$ |
|  |  | 2,690 |  |  |  |  |
|  |  | 1,371 |  |  |  |  |
|  |  | 1,614 |  |  |  |  |
|  |  | 1,494 |  |  |  |  |
| | KO | 1,929 | 9,494 | $0.0271 \pm 0.0012$ | $706 \pm 12$ | $82.9 \pm 1.2$ |
|  |  | 1,991 |  |  |  |  |
|  |  | 2,212 |  |  |  |  |
|  |  | 1,836 |  |  |  |  |
|  |  | 1,526 |  |  |  |  |
| 2 | WT | 1,652 | 9,093 | $0.0403 \pm 0.0046$ | $926 \pm 66$ | $93.1 \pm 5.7$ |
|  |  | 2,550 |  |  |  |  |
|  |  | 1,055 |  |  |  |  |
|  |  | 1,783 |  |  |  |  |
|  |  | 2,053 |  |  |  |  |
| | KO | 2,134 | 10,621 | $0.0469 \pm 0.0008$ | $838 \pm 16$ | $86.2 \pm 1.8$ |
|  |  | 2,218 |  |  |  |  |
|  |  | 2,512 |  |  |  |  |
|  |  | 1,971 |  |  |  |  |
|  |  | 1,786 |  |  |  |  |

Data are medians ± median absolute deviation (MAD). Five repetitions were carried out for each condition. WT, wild-type; KO, vimentin knock-out

**Table S3.** Force-mode atomic force microscopy analysis of dendritic cell mechanics.

| Extend | Cell type | Young's modulus |  | Relaxation time |  |
| --- | --- | --- | --- | --- | --- |
|  |  | Cells (n) | (Pa) | Cells (n) | (s) |
| 1 | WT | 12 | 826 ±575 | 8 | 1.7 ±1.1 |
| | KO | 16 | 280 ±248 | 13 | 1.9 ±1.8 |
| 2 | WT | 12 | 1,573 ±1,221 | 7 | 2.6 ±1.2 |
| | KO | 16 | 878 ±1,073 | 14 | 2.4 ±1.3 |
| 3 | WT | 11 | 2,617 ±2,295 | 8 | 3.1 ±1.6 |
| | KO | 16 | 1,118 ±993 | 17 | 3.4 ±1.5 |
| 4 | WT | 10 | 3,782 ±2,731 | 7 | 5.7 ±2.3 |
| | KO | 16 | 1,491 ±991 | 17 | 4.9 ±2.4 |
| 5 | WT | 9 | 10,275 ±7,669 | 6 | 4.9 ±1.7 |
| | KO | 16 | 3,288 ±2,583 | 17 | 4.8 ±1.0 |

Data are means ± standard deviation (SD); WT, wild-type; KO, vimentin knock-out

## References

Bénichou, O., C. Loverdo, M. Moreau, and R. Voituriez. 2011. Intermittent search strategies. Review of Modern Physics. 83:81–129.

Brown, M.J., J.A. Hallam, E. Colucci-Guyon, and S. Shaw. 2001. Rigidity of circulating lymphocytes is primarily conferred by vimentin intermediate filaments. J Immunol. 166:6640–6646.

Burridge, K., and C. Guilluy. 2016. Focal adhesions, stress fibers and mechanical tension. Exp Cell Res. 343:14–20.

Chabaud, M., M.L. Heuze, M. Bretou, P. Vargas, P. Maiuri, P. Solanes, M. Maurin, E. Terriac, M. Le Berre, D. Lankar, T. Piolot, R.S. Adelstein, Y. Zhang, M. Sixt, J. Jacobelli, O. Benichou, R. Voituriez, M. Piel, and A.M. Lennon-Dumenil. 2015. Cell migration and antigen capture are antagonistic processes coupled by myosin II in dendritic cells. Nat Commun. 6:7526.

Chan, C.J., A.E. Ekpenyong, S. Golfier, W. Li, K.J. Chalut, O. Otto, J. Elgeti, J. Guck, and F. Lautenschläger. 2015. Myosin II activity softens cells in suspension. Biophysical journal. 108:1856–1869.

Chen, J.-T. 2005. Using the sum-of-uniforms method to generate correlated random variates with certain marginal distribution. European Journal of Operational Research. 167:226–242.

Chernoivanenko, I.S., A.A. Minin, and A.A. Minin. 2013. Role of vimentin in cell migration. Ontogenez. 44:186–202.

Chugh, P., A.G. Clark, M.B. Smith, D.A.D. Cassani, K. Dierkes, A. Ragab, P.P. Roux, G. Charras, G. Salbreux, and E.K. Paluch. 2017. Actin cortex architecture regulates cell surface tension. Nat Cell Biol. 19:689–697.

Costigliola, N., L. Ding, C.J. Burckhardt, S.J. Han, E. Gutierrez, A. Mota, A. Groisman, T.J. Mitchison, and G. Danuser. 2017. Vimentin fibers orient traction stress. Proc Natl Acad Sci USA 114:5195–5200.

Danielsson, F., M.K. Peterson, H. Caldeira Araujo, F. Lautenschläger, and A.K.B. Gad. 2018. Vimentin diversity in health and disease. Cells. 7:147.

De Pascalis, C., C. Pérez-González, S. Seetharaman, B. Boëda, B. Vianay, M. Burute, C. Leduc, N. Borghi, X. Trepat, and S. Etienne-Manneville. 2018. Intermediate filaments control collective migration by restricting traction forces and sustaining cell-cell contacts. J Cell Biol. 217:3031–3044.

Eriksson, J.E., T. He, A.V. Trejo-Skalli, A.S. Härmälä-Braskén, J. Hellman, Y.H. Chou, and R.D. Goldman. 2004. Specific in vivo phosphorylation sites determine the assembly dynamics of vimentin intermediate filaments. J Cell Sci. 117:919–932.

Faure-Andre, G., P. Vargas, M.I. Yuseff, M. Heuze, J. Diaz, D. Lankar, V. Steri, J. Manry, S. Hugues, F. Vascotto, J. Boulanger, G. Raposo, M.R. Bono, M. Rosemblatt, M. Piel, and A.M. Lennon-Dumenil. 2008. Regulation of dendritic cell migration by CD74, the MHC class II-associated invariant chain. Science. 322:1705–1710.

Fischer-Friedrich, E., Y. Toyoda, C.J. Cattin, D.J. Müller, A.A. Hyman, and F. Jülicher. 2016. Rheology of the Active Cell Cortex in Mitosis. Biophysical journal. 111:589–600.

Fletcher, D.A., and R.D. Mullins. 2010. Cell mechanics and the cytoskeleton. Nature. 463:485–492.

Gan, Z., L. Ding, Christoph J. Burckhardt, J. Lowery, A. Zaritsky, K. Sitterley, A. Mota, N. Costigliola, Colby G. Starker, Daniel F. Voytas, J. Tytell, Robert D. Goldman, and G. Danuser. 2016. Vimentin intermediate filaments template microtubule networks to enhance persistence in cell polarity and directed migration. Cell Syst. 3:252–263.e258.

Guo, M., A.J. Ehrlicher, S. Mahammad, H. Fabich, M.H. Jensen, J.R. Moore, J.J. Fredberg, R.D. Goldman, and D.A. Weitz. 2013. The role of vimentin intermediate filaments in cortical and cytoplasmic mechanics. Biophysical journal. 105:1562–1568.

Hampton, H.R., and T. Chtanova. 2019. Lymphatic migration of immune cells. Frontiers in immunology. 10:1168.

Hawkins, R.J., M. Piel, G. Faure-Andre, A.M. Lennon-Dumenil, J.F. Joanny, J. Prost, and R. Voituriez. 2009. Pushing off the walls: a mechanism of cell motility in confinement. Phys Rev Lett. 102:058103.

Helfand, B.T., M.G. Mendez, S.N.P. Murthy, D.K. Shumaker, B. Grin, S. Mahammad, U. Aebi, T. Wedig, Y.I. Wu, K.M. Hahn, M. Inagaki, H. Herrmann, and R.D. Goldman. 2011. Vimentin organization modulates the formation of lamellipodia. Mol Biol Cell. 22:1274–1289.

Herbig, M., A. Mietke, P. Muller, and O. Otto. 2018. Statistics for real-time deformability cytometry: clustering, dimensionality reduction, and significance testing. Biomicrofluidics. 12:042214.

Heuzé, M.L., O. Collin, E. Terriac, A.M. Lennon-Dumenil, and M. Piel. 2011. Cell migration in confinement: a micro-channel-based assay. Methods Mol Biol. 769:415–434.

Heuzé, M.L., P. Vargas, M. Chabaud, M. Le Berre, Y.J. Liu, O. Collin, P. Solanes, R. Voituriez, M. Piel, and A.M. Lennon-Dumenil. 2013. Migration of dendritic cells: physical principles, molecular mechanisms, and functional implications. Immunological reviews. 256:240–254.

Hu, J., Y. Li, Y. Hao, T. Zheng, S.K. Gupta, G.A. Parada, H. Wu, S. Lin, S. Wang, X. Zhao, R.D. Goldman, S. Cai, and M. Guo. 2019. High stretchability, strength, and toughness of living cells enabled by hyperelastic vimentin intermediate filaments. Proc Natl Acad Sci USA. 116:17175–17180.

Ivaska, J., H.M. Pallari, J. Nevo, and J.E. Eriksson. 2007. Novel functions of vimentin in cell adhesion, migration, and signaling. Exp Cell Res. 313:2050–2062.

Janmey, P.A., U. Euteneuer, P. Traub, and M. Schliwa. 1991. Viscoelastic properties of vimentin compared with other filamentous biopolymer networks. J Cell Biol. 113:155–160.

Jiu, Y., J. Lehtimaki, S. Tojkander, F. Cheng, H. Jaalinoja, X. Liu, M. Varjosalo, J.E. Eriksson, and P. Lappalainen. 2015. Bidirectional interplay between vimentin intermediate filaments and contractile actin stress fibers. Cell Rep. 11:1511–1518.

Jiu, Y., J. Peranen, N. Schaible, F. Cheng, J.E. Eriksson, R. Krishnan, and P. Lappalainen. 2017. Vimentin intermediate filaments control actin stress fiber assembly through GEF-H1 and RhoA. J Cell Sci. 130:892–902.

Kopf, A., J. Renkawitz, R. Hauschild, I. Girkontaite, K. Tedford, J. Merrin, O. Thorn-Seshold, D. Trauner, H. Häcker, K.-D. Fischer, E. Kiermaier, and M. Sixt. 2020. Microtubules control cellular shape and coherence in amoeboid migrating cells. J Cell Biol. 219:e201907154.

Korb, T., K. Schluter, A. Enns, H.U. Spiegel, N. Senninger, G.L. Nicolson, and J. Haier. 2004. Integrity of actin fibers and microtubules influences metastatic tumor cell adhesion. Exp Cell Res. 299:236–247.

Lakhan, V.C. 1981. Generating autocorrelated pseudo-random numbers with specific distributions. Journal of Statistical Computation and Simulation. 12:303–309.

Lämmermann, T., B.L. Bader, S.J. Monkley, T. Worbs, R. Wedlich-Soldner, K. Hirsch, M. Keller, R. Forster, D.R. Critchley, R. Fassler, and M. Sixt. 2008. Rapid leukocyte migration by integrin-independent flowing and squeezing. Nature. 453:51–55.

Lämmermann, T., J. Renkawitz, X. Wu, K. Hirsch, C. Brakebusch, and M. Sixt. 2009. Cdc42-dependent leading edge coordination is essential for interstitial dendritic cell migration. Blood. 113:5703–5710.

Lauffenburger, D.A., and A.F. Horwitz. 1996. Cell migration: a physically integrated molecular process. Cell. 84:359–369.

Lautenschläger, F., S. Paschke, S. Schinkinger, A. Bruel, M. Beil, and J. Guck. 2009. The regulatory role of cell mechanics for migration of differentiating myeloid cells. Proc Natl Acad Sci U S A. 106:15696–15701.

Lautenschläger, F., and M. Piel. 2013. Microfabricated devices for cell biology: all for one and one for all. Curr Opin Cell Biol. 25:116–124.

Le Berre, M., E. Zlotek-Zlotkiewicz, D. Bonazzi, F. Lautenschlaeger, and M. Piel. 2014. Methods for two-dimensional cell confinement. Methods in cell biology. 121:213–229.

Liu, Y.J., M. Le Berre, F. Lautenschläger, P. Maiuri, A. Callan-Jones, M. Heuze, T. Takaki, R. Voituriez, and M. Piel. 2015. Confinement and low adhesion induce fast amoeboid migration of slow mesenchymal cells. Cell. 160:659–672.

M. Reza Shaebani, R.J., Ludger Santen, Luiza Stankevicins, and Franziska Lautenschläger. 2020. Persistence-speed coupling enhances the search efficiency of migrating immune cells. arXiv.

Maiuri, P., J.F. Rupprecht, S. Wieser, V. Ruprecht, O. Benichou, N. Carpi, M. Coppey, S. De Beco, N. Gov, C.P. Heisenberg, C. Lage Crespo, F. Lautenschläger, M. Le Berre, A.M. Lennon-Dumenil, M. Raab, H.R. Thiam, M. Piel, M. Sixt, and R. Voituriez. 2015. Actin flows mediate a universal coupling between cell speed and cell persistence. Cell. 161:374–386.

Maiuri, P., E. Terriac, P. Paul-Gilloteaux, T. Vignaud, K. McNally, J. Onuffer, K. Thorn, P.A. Nguyen, N. Georgoulia, D. Soong, A. Jayo, N. Beil, J. Beneke, J.C. Lim, C.P. Sim, Y.S. Chu, W.C.R. participants, A. Jimenez-Dalmaroni, J.F. Joanny, J.P. Thiery, H. Erfle, M. Parsons, T.J. Mitchison, W.A. Lim, A.M. Lennon-Dumenil, M. Piel, and M. Thery. 2012. The first world cell race. Current biology: CB. 22:R673–675.

Mendez, M.G., S.-I. Kojima, and R.D. Goldman. 2010. Vimentin induces changes in cell shape, motility, and adhesion during the epithelial to mesenchymal transition. FASEB J. 24:1838–1851.

Mendez, M.G., D. Restle, and P.A. Janmey. 2014. Vimentin enhances cell elastic behavior and protects against compressive stress. Biophysical journal. 107:314–323.

Mokbel, M., D. Mokbel, A. Mietke, N. Träber, S. Girardo, O. Otto, J. Guck, and S. Aland. 2017. Numerical simulation of real-time deformability cytometry to extract cell mechanical properties. ACS Biomater Sci Eng. 3:2962–2973.

Moreau, H.D., C. Blanch-Mercader, R. Attia, M. Maurin, Z. Alraies, D. Sanséau, O. Malbec, M.-G. Delgado, P. Bousso, J.-F. Joanny, R. Voituriez, M. Piel, and A.-M. Lennon-Duménil. 2019. Macropinocytosis overcomes directional bias in dendritic cells due to hydraulic resistance and facilitates space exploration. Dev Cell. 49:171–188.e175.

Ng, L.G., A. Hsu, M.A. Mandell, B. Roediger, C. Hoeller, P. Mrass, A. Iparraguirre, L.L. Cavanagh, J.A. Triccas, S.M. Beverley, P. Scott, and W. Weninger. 2008. Migratory dermal dendritic cells act as rapid sensors of protozoan parasites. PLoS Pathog. 4:e1000222.

Ngan, C.Y., H. Yamamoto, I. Seshimo, T. Tsujino, M. Man-i, J.I. Ikeda, K. Konishi, I. Takemasa, M. Ikeda, M. Sekimoto, N. Matsuura, and M. Monden. 2007. Quantitative evaluation of vimentin expression in tumour stroma of colorectal cancer. Br J Cancer. 96:986–992.

Nieminen, M., T. Henttinen, M. Merinen, F. Marttila-Ichihara, J.E. Eriksson, and S. Jalkanen. 2006. Vimentin function in lymphocyte adhesion and transcellular migration. Nat Cell Biol. 8:156–162.

Otto, O., P. Rosendahl, A. Mietke, S. Golfier, C. Herold, D. Klaue, S. Girardo, S. Pagliara, A. Ekpenyong, A. Jacobi, M. Wobus, N. Topfner, U.F. Keyser, J. Mansfeld, E. Fischer-Friedrich, and J. Guck. 2015. Real-time deformability cytometry: on-the-fly cell mechanical phenotyping. Nature methods. 12:199–202.

Patteson, A.E., K. Pogoda, F.J. Byfield, K. Mandal, Z. Ostrowska-Podhorodecka, E.E. Charrier, P.A. Galie, P. Deptuła, R. Bucki, C.A. McCulloch, and P.A. Janmey. 2019a. Loss of vimentin enhances cell motility through small confining spaces. Small. 15:1903180.

Patteson, A.E., A. Vahabikashi, K. Pogoda, S.A. Adam, K. Mandal, M. Kittisopikul, S. Sivagurunathan, A. Goldman, R.D. Goldman, and P.A. Janmey. 2019b. Vimentin protects cells against nuclear rupture and DNA damage during migration. J Cell Biol. 218:4079–4092.

Randolph, G.J., V. Angeli, and M.A. Swartz. 2005. Dendritic-cell trafficking to lymph nodes through lymphatic vessels. Nature reviews. Immunology. 5:617–628.

Renkawitz, J., A. Kopf, J. Stopp, I. de Vries, M.K. Driscoll, J. Merrin, R. Hauschild, E.S. Welf, G. Danuser, R. Fiolka, and M. Sixt. 2019. Nuclear positioning facilitates amoeboid migration along the path of least resistance. Nature. 568:546–550.

Reversat, A., F. Gaertner, J. Merrin, J. Stopp, S. Tasciyan, J. Aguilera, I. de Vries, R. Hauschild, M. Hons, M. Piel, A. Callan-Jones, R. Voituriez, and M. Sixt. 2020. Cellular locomotion using environmental topography. Nature. 582:582–585.

Sarria, A.J., J.G. Lieber, S.K. Nordeen, and R.M. Evans. 1994. The presence or absence of a vimentin-type intermediate filament network affects the shape of the nucleus in human SW-13 cells. J Cell Sci. 107:1593–1607.

Satelli, A., and S. Li. 2011. Vimentin in cancer and its potential as a molecular target for cancer therapy. Cell Mol Life Sci. 68:3033–3046.

Schindelin, J., I. Arganda-Carreras, E. Frise, V. Kaynig, M. Longair, T. Pietzsch, S. Preibisch, C. Rueden, S. Saalfeld, B. Schmid, J.Y. Tinevez, D.J. White, V. Hartenstein, K. Eliceiri, P. Tomancak, and A. Cardona. 2012. Fiji: an open-source platform for biological-image analysis. Nature methods. 9:676–682.

Schon-Hegrad, M.A., J. Oliver, P.G. McMenamin, and P.G. Holt. 1991. Studies on the density, distribution, and surface phenotype of intraepithelial class II major histocompatibility complex antigen (Ia)-bearing dendritic cells (DC) in the conducting airways. J Exp Med. 173:1345–1356.

Smoler, M., G. Coceano, I. Testa, L. Bruno, and V. Levi. 2020. Apparent stiffness of vimentin intermediate filaments in living cells and its relation with other cytoskeletal polymers. Biochim Biophys Acta Mol Cell Res. 1867:118726.

Stankevicins, L., N. Ecker, E. Terriac, P. Maiuri, R. Schoppmeyer, P. Vargas, A.-M. Lennon-Duménil, M. Piel, B. Qu, M. Hoth, K. Kruse, and F. Lautenschläger. 2020. Deterministic actin waves as generators of cell polarization cues. Proc Natl Acad Sci U S A 117:826–835.

Tejedor, V., R. Voituriez, and O. Bénichou. 2012. Optimizing persistent random searches. Phys Rev Lett. 108:088103.

Urbanska, M., P. Rosendahl, M. Krater, and J. Guck. 2018. High-throughput single-cell mechanical phenotyping with real-time deformability cytometry. Methods in cell biology. 147:175–198.

Vargas, P., P. Maiuri, M. Bretou, P.J. Saez, P. Pierobon, M. Maurin, M. Chabaud, D. Lankar, D. Obino, E. Terriac, M. Raab, H.R. Thiam, T. Brocker, S.M. Kitchen-Goosen, A.S. Alberts, P. Sunareni, S. Xia, R. Li, R. Voituriez, M. Piel, and A.M. Lennon-Dumenil. 2016. Innate control of actin nucleation determines two distinct migration behaviours in dendritic cells. Nat Cell Biol. 18:43–53.

Vargas, P., E. Terriac, A.M. Lennon-Dumenil, and M. Piel. 2014. Study of cell migration in microfabricated channels. Journal of visualized experiments: JoVE:e51099.

Willemain, R.T., and A.P. Desautels. 1993. A method to generate autocorrelated uniform random numbers. Journal of Statistical Computation and Simulation. 45:23–31.

Wu, P.H., A. Giri, S.X. Sun, and D. Wirtz. 2014. Three-dimensional cell migration does not follow a random walk. Proc Natl Acad Sci U S A 111:3949–3954.

